# *Myxococcus xanthus*, a nonpathogenic bacterium, eliminates *Cryptococcus neoformans*, a fungal pathogen of human, independent of cell-cell contact

**DOI:** 10.1101/2022.08.25.505288

**Authors:** Huan Zhang, Joshua D. Pettibon, Raj Patel, Beiyan Nan

**Affiliations:** Department of Biology, Texas A&M University, College Station, TX77845, USA

**Keywords:** antifungal, fungicidal, bacteria-fungal interaction, cell envelop integrity, microbial ecology

## Abstract

Each year an estimated 1.2 billion people suffer from fungal diseases and 1.5 - 2 million die from fungal infections - surpassing the loss from malaria and tuberculosis^1-3^. Due to the similarities between fungal and human cells, the lack of fungal-specific targets has become the major hurdle for antifungal discovery. Many fungi, including the deadly human pathogen *Cryptococcus neoformans*, are found in soil, where they compete with other microorganisms, including bacteria. However, most bacteria that inhibit fungal growth are pathogens and their antifungal effects strictly rely on cell-cell contact. Here we show that *Myxococcus xanthus*, a nonpathogenic, soil-dwelling bacterium, efficiently eliminates *C. neoformans* and strongly inhibits the production of fungal virulence factors. Remarkably, these antifungal activities do not require cell-cell contact. Using fluorescence microscopy, we found that *M. xanthus* increases the permeability of *C. neoformans* cells. Our results on the cross-kingdom interaction between *M. xanthus* and *C. neoformans* will reveal fundamental mechanisms for bacterial-fungal interactions and suggest novel strategies for antifungal therapies.

## Introduction

Invasive fungal diseases impose a major public health burden, with mortality rates between 30 – 90%^1-4^. About 90% of the deaths due to invasive fungal infections are caused by *Cryptococcus, Aspergillus, Candida*, and *Pneumocystis*^1,5^. *Cryptococcus* species are among the most significant threats to human health and the burden of cryptococcal meningitis alone exceeds 600,000 cases annually^6,7^. Ubiquitous and fully virulent in the environment, *Cryptococcus* are commonly isolated from soil and avian inhabitats^8,9^. Humans are exposed to *Cryptococcus* through the inhalation of spores and desiccated yeast cells, but such exposure is typically asymptomatic. Serological surveys show that over 80% of children in urban environments have been infected^10^. However, for immunocompromised individuals owing to HIV infection, cancer, organ transplant, and COVID-19 treatments, *Cryptococcus* can disseminate from the lungs to the bloodstream and infect any tissue^11-17^. Remarkably, *Cryptococcus* are among a few microorganisms that can infiltrate the blood-brain barrier and infect the central nervous system, which is fatal without effective treatments^7,18,19^. In recent years, Cryptococcosis outbreaks have become more common in North America, including in British Columbia, and in Washington and Oregon in the US^1^. Effective methods to inhibit growth and infection of these pathogens are urgently needed to reduce mortality and improve patient care.

Despite the rapid evolving drug-resistance, drug development has stalled and no antifungals have been approved since 2006^20^. A major challenge to the development of antifungals is the lack of fungal-specific targets. Nevertheless, such targets, as well as antifungal mechanisms must have existed for millions of years in the environmental niches shared by fungi and bacteria. Researchers have realized the breadth of fungal-bacterial interactions and started to explore the possibility of controlling fungal growth using bacteria. Unfortunately, most of the reported antifungal mechanisms are not applicable to therapies. First, many bacteria that antagonize fungi, such as *Staphylococcus aureus, Pseudomonas aeruginosa, Serratia marcescens, Klebsiella pneumoniae, Streptococcus mitis*, etc., are pathogens themselves^21-26^. Second, most of the reported inhibitory interactions strictly require cell-cell contact between live fungi and bacteria^21,22,25,27^. Third, some bacteria, such as *P. aeruginosa, Escherichia coli, Clostridioides difficile*, and *S. aureus*, gain virulence and antibiotic resistance from their interactions with fungi^28-32^. Thus, antifungal activities from nonpathogenic bacteria that do not require cell-cell contact promise applicable strategies to combat pathogenic fungi.

The gram-negative myxobacteria are ubiquitous in soil^33,34^. Sharing common habitats, myxobacteria and *Cryptococcus* must have been competing and coevolving for millions of years. Myxobacteria distinguish themselves from other bacteria by their complex life cycles and social behaviors. For example, these bacteria can form multicellular fruiting bodies and cooperatively prey on other microorganisms^35,36^. Such complexity is matched by their extraordinary genomes of up to 16 Mbp^37^, which implies exceptional potential for secondary metabolite production and cell-cell communication. We found that *Myxococcus xanthus*, a soil-dwelling myxobacterium that shares habitats with *C. neoformans* in nature, eliminates *C. neoformans* efficiently, reduces the production of fungal virulence factors, melanin and capsule, and inhibits the formation of fungal biofilms. In contrast, *C. neoformans* does not inhibit the growth of *M. xanthus*. Remarkably, cell-cell contact is not essential for the fungicidal activities of *M. xanthus*. The cross-kingdom interaction between *M. xanthus* and *C. neoformans* may suggest new approaches to combat fungal infection and new targets for antifungal drugs.

## Results and Discussion

### *M. xanthus* efficiently eliminates *C. neoformans*

To study their potential interactions, we grew *C. neoformans* into a lawn and spotted the suspended cells of wild-type *M. xanthus* onto it. After 48 h of incubation, we found that *M. xanthus* eradicated *C. neoformans* at its inoculation sites (**Fig. 1A**). To test if this inhibition happens in aqueous environments and to quantify the inhibitory efficiency, *C. neoformans* and *M. xanthus* were grown separately in liquid culture and mixed to a ratio of biomass *C. neo* : *M. xan* of 3 : 1. After 48 h of incubation, the survival rates of both organisms were quantified on plates by colony-forming units (cfu). We found that *M. xanthus* reduced the cfu of *C. neoformans* by ∼10^4^-fold (**Fig. 1B**). Such a strong inhibitory effect qualifies as fungicidal (>10^3^-fold reduction)^38^. In contrast, the growth of *M. xanthus* was not affected by *C. neoformans* (**Fig. 1C**). The fungicidal activities of *M. xanthus* are not due to simple competition of resources in the medium, as *Escherichia coli* did not inhibit the growth of *C. neoformans* (**Fig. 1B**).

**Fig. 1.**
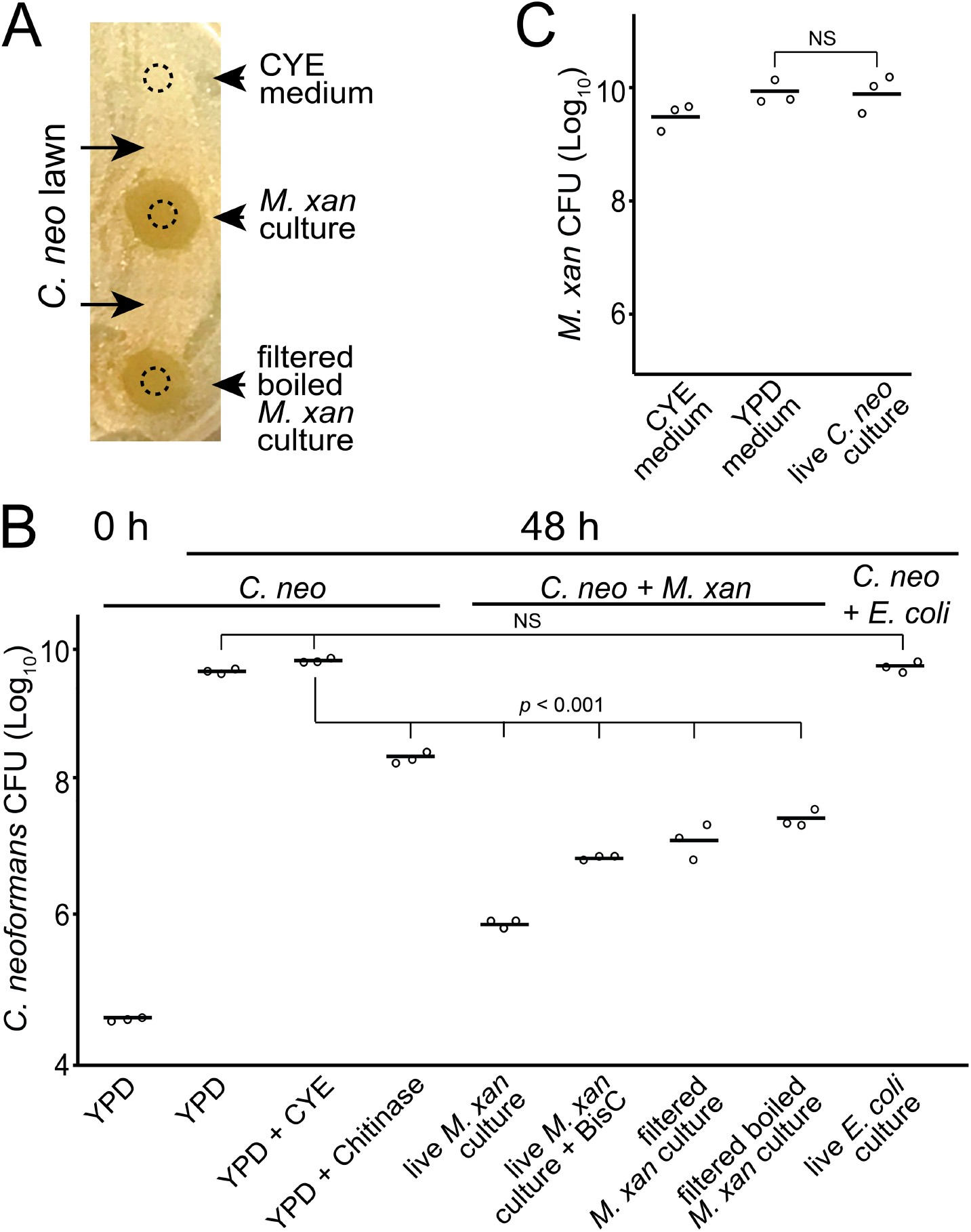
*M. xanthus* (*M. xan*) eliminates *C. neoformans* (*C. neo*) efficiently. **A)** Live and filtrated and boiled *M. xanthus* culture eliminate *C. neoformans* on solid medium. Dotted circles show the initial inoculation sites. **B)** The killing efficiency was quantified in liquid culture. **C)** *C. neoformans* does not inhibit the growth of *M. xanthus*. NS, nonsignificant. *p* values were calculated using the Student paired t test with a two-tailed distribution here and in all subsequent figures. BisC, bisdionine.

### Cell-cell contact enhances, but is not required for, the fungicidal activities of *M. xanthus*

To test if the *M. xanthus* fungicidal activities rely on cell-cell contact, we performed a two-chamber assay. Liquid *M. xanthus* culture or blank medium were inoculated into the bottom chambers of a 24-well plate that contains 0.4-μm pore size membrane inserts. *C. neoformans* was inoculated on the top chamber (**Fig. 2A**). Even when separated by the membrane, *M. xanthus* still efficiently eliminated *C. neoformans* cells, by ∼10^3^-fold. Thus, though cell-cell contact may enhance the killing efficiency, the major fungicidal factors are in the supernatant fraction of *M. xanthus* culture (**Fig. 2B**). To further test the nature of such factors, we passed the supernatant of *M. xanthus* monoculture through 0.2-μm pores and the filtered medium still killed *C. neoformans*. Importantly, additional boiling of the supernatant did not reduce the killing efficiency further, indicating that the fungicidal factors of *M. xanthus* are heat-resistant (**Fig. 1B, 2B**). The heat-stability of the fungicidal factors suggests that they may mainly consist of ions, small metabolites or short peptides rather than large proteins.

**Fig. 2.**
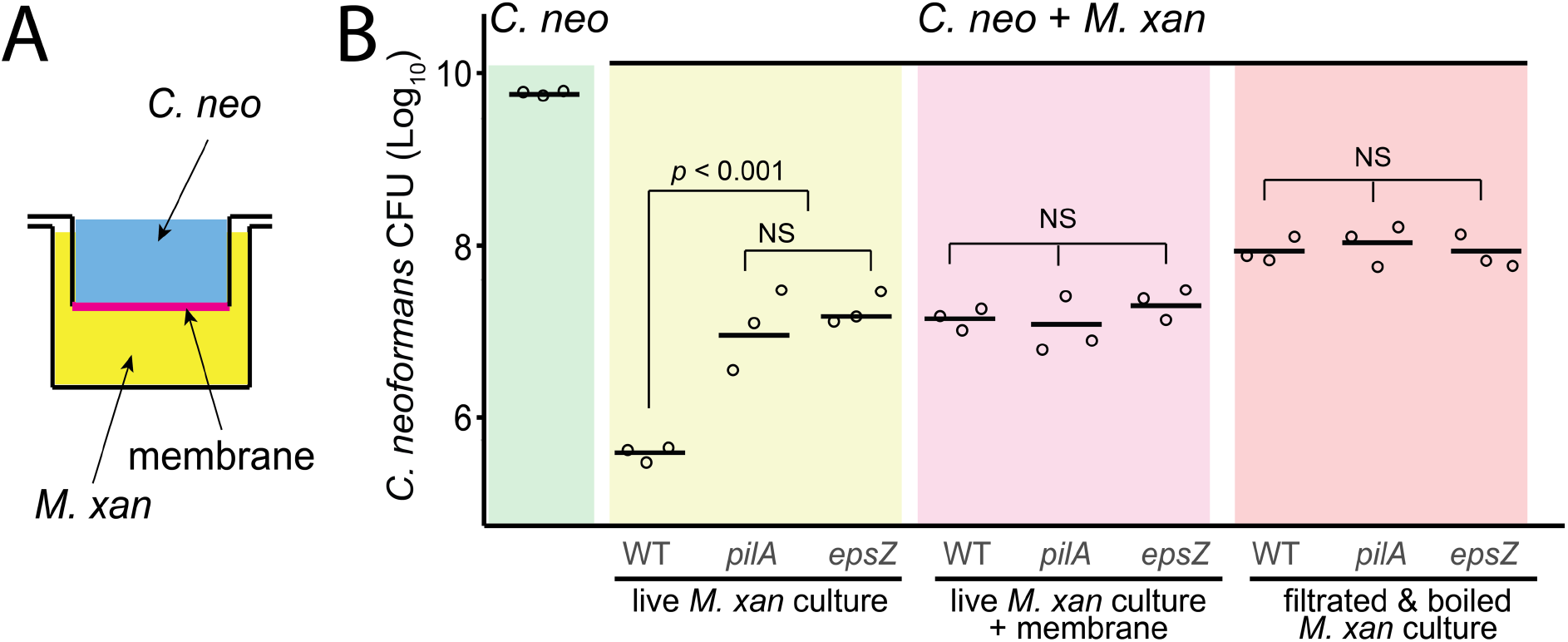
Cell-cell contact is not essential for the fungicidal activities of *M. xanthus*. **A)** The experiment set up. **B)** *M. xan* still kills *C. neoformans* when the two organisms are separated by a membrane. Two *M. xanthus* mutants, *pilA* and *epsZ*, both are less likely to make contacts with *C. neoformans*, eliminate the fungus with wild-type like efficiencies in and membrane separated coculture.

What is the role of cell-cell contact in the warfare between *M. xanthus* and *C. neoformans*? *M. xanthus* cells may form intra- and interspecies cell-cell contact through type IV pili and extracellular polysaccharides (**EPS**)^39-41^. We tested the killing efficiency of the *M. xanthus pilA*^42^ and *epsZ*^43^ strains that are unable to produce pilus and EPS, respectively, and thus are less likely to make direct contact with *C. neoformans*. We found that both mutants showed reduced killing abilities against *C. neoformans*. Strikingly, in the presence of membranes, their killing efficiencies were similar to the wild type cells (**Fig. 2B**). Taken together, these results indicate that rather than mediating the killing directly, cell-cell contact may enhance killing efficiency by increasing the local concentrations of the fungicidal factors. Thus, in contrast to other antifungal bacteria such as *P. aeruginosa*^21^ and *Bacillus safensis*^27^, cell-cell contact is not required for the fungicidal effects of *M. xanthus*.

### *M. xanthus* inhibits the production of *C. neoformans* virulence factors

To investigate if *M. xanthus* affects the virulence of *C. neoformans*, we quantified the production of the two major cryptococcal virulence factors, melanin and capsule. As melanin is not readily detectable on regular YPD medium, we induced its production using L-3,4-dihydroxyphenyl-alanine (**L-DOPA**). Compared to the pure *C. neoformans* culture, the presence of *M. xanthus* abolished melanin production (**Fig. 3A**). One may argue that the reduced production of melanin is due to the loss of viable fungal cells. To test if *M. xanthus* reduces the virulence of individual *C. neoformans* cells, we used Dulbecco’s Modified Eagle Medium (**DMEM**) to induce capsule formation at 37 °C. Although 37 °C is not its optimum temperature (30 - 32 °C), *M. xanthus* still reduced the thickness of individual *C. neoformans* capsules by 70% (**Fig. 3A, 3B**). As capsular polysaccharides are required for biofilm formation, which is linked to the pathogenicity of *C. neoformans*, we quantified biofilm formation using crystal violet that binds to EPS. The *M. xanthus pilA* strain that does not form biofilms itself significantly reduced the binding of crystal violet in the co-culture (**Fig. 3C**). Taken together, besides killing *C. neoformans* directly, *M. xanthus* also reduces the virulence of this fungal pathogen.

**Fig. 3.**
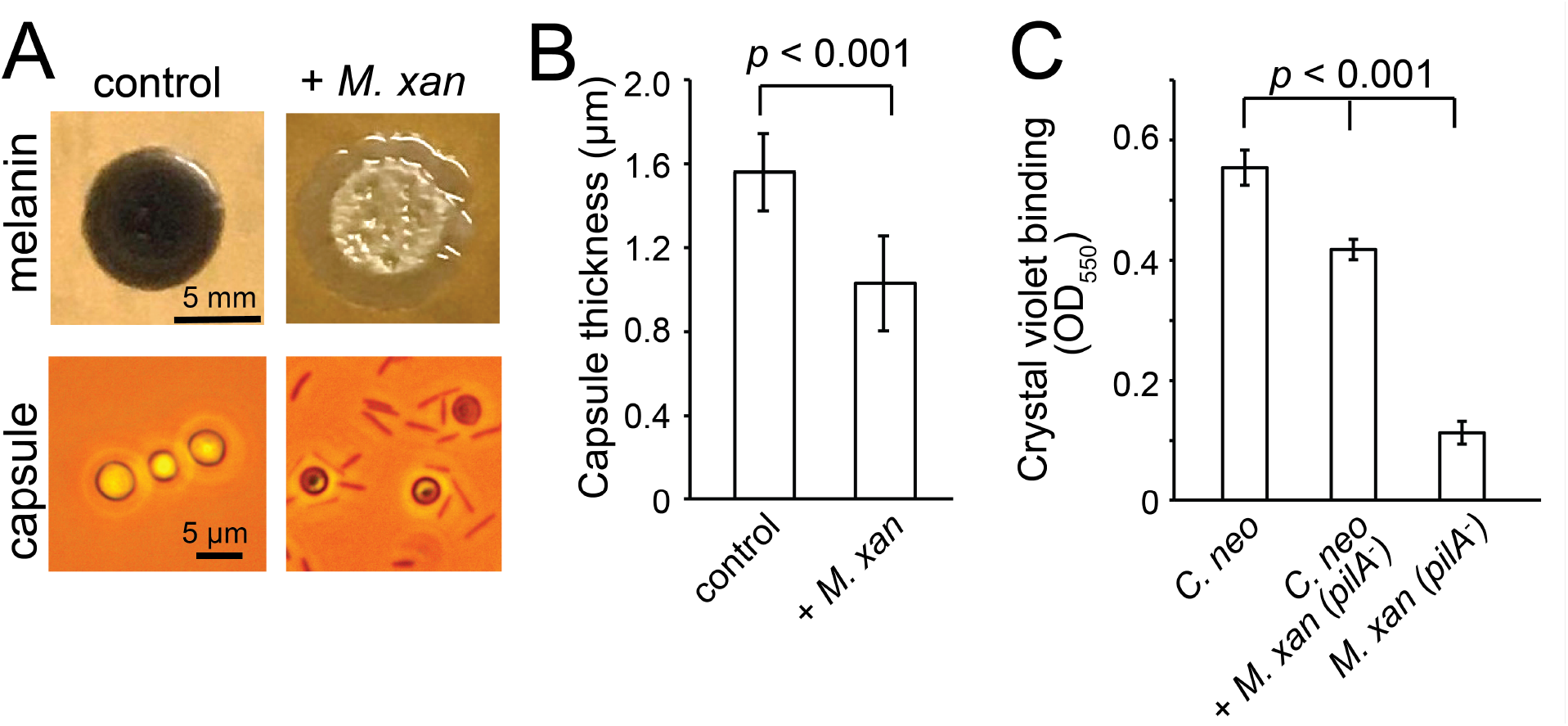
*M. xanthus* inhibits the production of *C. neoformans* virulence factors melanin and capsule and reduces the formation of fungal biofilms. **A)** Melanin and capsule production of *C. neoformans* on L-DOPA and DMEM plate, respectively. **B)** Quantitative analysis of capsule thickness. **C)** Quantitative analysis of biofilm formation using the absorption of crystal violet at 550 nm. Error bars show standard derivations here and in all subsequent figures.

### *M. xanthus* disrupts the integrity of the *C. neoformans* cell envelope

To visualize the fungicidal effects of *M. xanthus*, we investigated the morphology of *C. neoformans* cells within 6 h of contact with *M. xanthus*, before mass cell death occurs. Compared to the cells in monoculture, when grown together with *M. xanthus, C. neoformans* showed significant disorders in the cytoplasm (**Fig. 4A**). We hypothesized that such disorders could result from the degradation of cell envelopes. To test this hypothesis, we investigated the integrity of both the *C. neoformans* cell wall and membrane using fluorescence microscopy. We used the chitin-binding dye calcofluor white (**CWF**) to stain chitin polymers in healthy cell wall and FM4-64-labeled wheat germ agglutinin (**WGA**) to detect chito-oligomers from cell wall degradation. In contrast to the fungal cells in monoculture that only showed weak binding to WGA, *C. neoformans* in the dual-species culture were heavily stained in the cytoplasm by WGA, and the foci of WGA co-localized with the disordered structures in bright-field images (**Fig. 4A**). We then tested the permeability of *C. neoformans* membrane using two fluorescent reporters, Bis-(1,3-dibutylbarbituric acid) trimethine oxonol (**DiBAC**_**4**_**(3)**)^44^ and propidium iodine (**PI**)^45^. Though minor damage, such as the depolarization of membranes can cause the internalization of DiBAC_4_(3), PI in cytoplasm often indicates severe membrane fracture, such as the formation of pores^22^. Neither dyes are able to enter the cytoplasm of *C. neoformans* in monocultures (**Fig. 4B**). In contrast, the fungal cells accumulated both DiBAC_4_(3) and PI significantly when co-cultivated with *M. xanthus* (**Fig. 4B**). To confirmed the severity of the membrane damage, we quantified the internalization of PI in the *C. neoformans* cells treated by amphotericin B (which induces membrane depolarization without pore formation, 3 μg/ml, for 6 h) and mellitin (also known as melittin, a pore-forming peptide, 10 μM, for 10 min)^22^. Our data indicated that mellitin, but not amphotericin B, resulted in the accumulation of PI in the fungal cytoplasm (**Fig. 4C**). Taken together, *M. xanthus* inflicts severe damages to both the *C. neoformans* cell wall and membranes. Some bacteria, such as *B. safensis*, uses chitinases as major antifungal factors^27^. Consistent with a prior report^27^, chitinases (100 µg/ml) showed limited inhibition on *C. neoformans* growth in monoculture. Different from *B. safensis, M. xanthus* still eliminated the fungus efficiently in the presence of the chitinase inhibitor bisdionine (**BisC**, 2.5 mM, **Fig. 1B**). Thus, Chitinases are heat-sensitive antifungal facctors that play secondary roles in the fungicidal activities of *M. xanthus*.

**Fig. 4.**
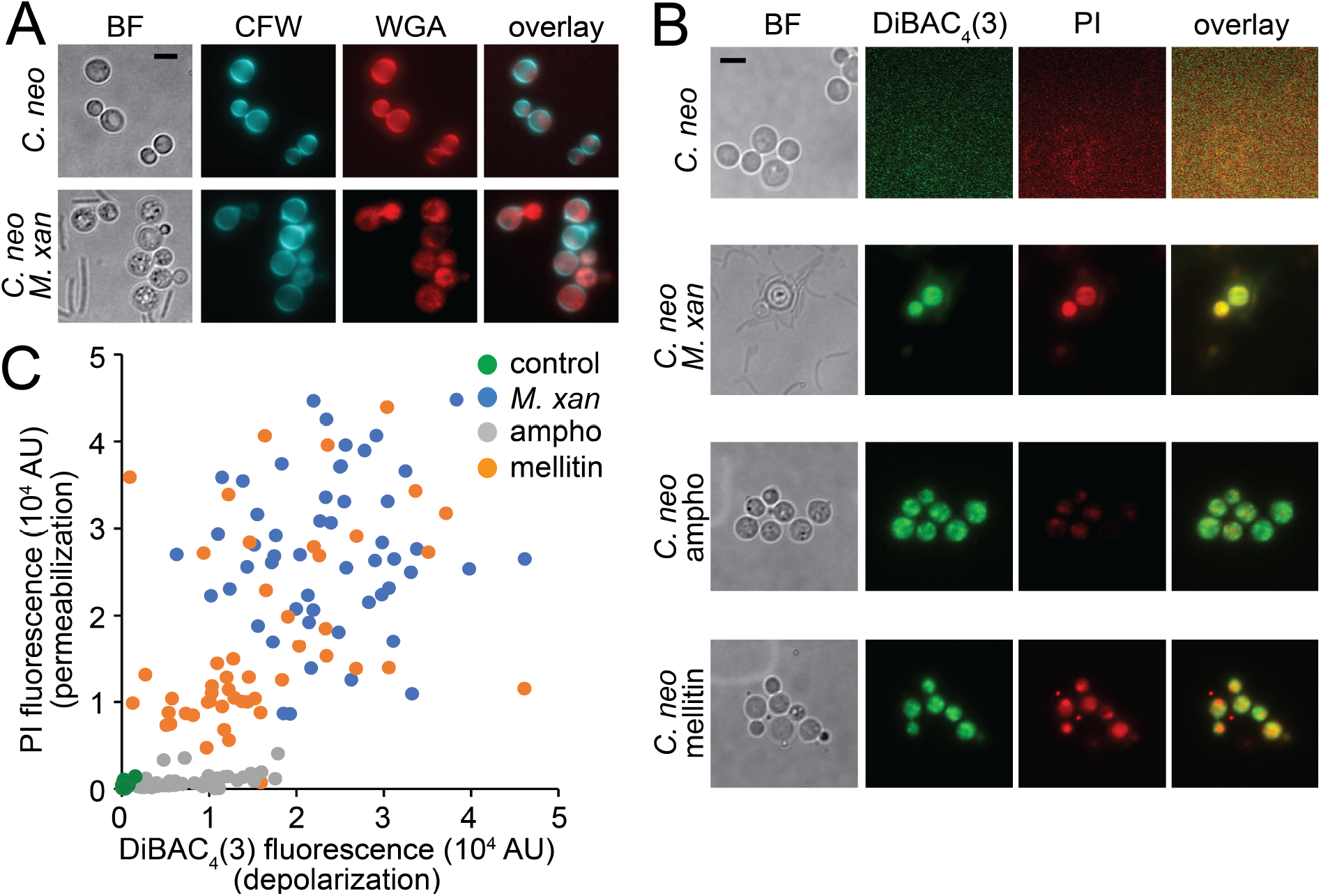
*M. xanthus* disrupts the integrity of *C. neoformans* cell envelope. **A)** *M. xanthus* causes the internalization of chito-oligomers in *C. neformanso* cells, which indicates the degradation of cell wall. **B)** *M. xanthus* increases the permeability of DiBAC_4_(3) and PI across *C. neoformans* membranes. The permeability of PI especially indicates the formation of pores in fungal membrane. Amphotericin B (ampho) and mellitin were used to depolarize and permeabilize membranes, respectively. **C)** The internalization of DiBAC_4_(3) and PI in 50 cells under each condition. Scale bars, 5 µm.

### *M. xanthus* shows potential broad-spectrum antifungal activities

*M. xanthus* also significantly inhibited the growth of *C. albicans*, another major fungal pathogen for humans (**Fig. 5A, 5B**). Different from *C. neoformans*, hypha is the virulent form of *C. albicans*^46^. As *M. xanthus* strongly inhibits the yeast-to-hyphal transition *of C. albicans*, it reduces the virulence of this pathogen (**Fig. 5C**). Thus, our data indicate that *M. xanthus* has broad-spectrum antifungal activities.

**Fig. 5.**
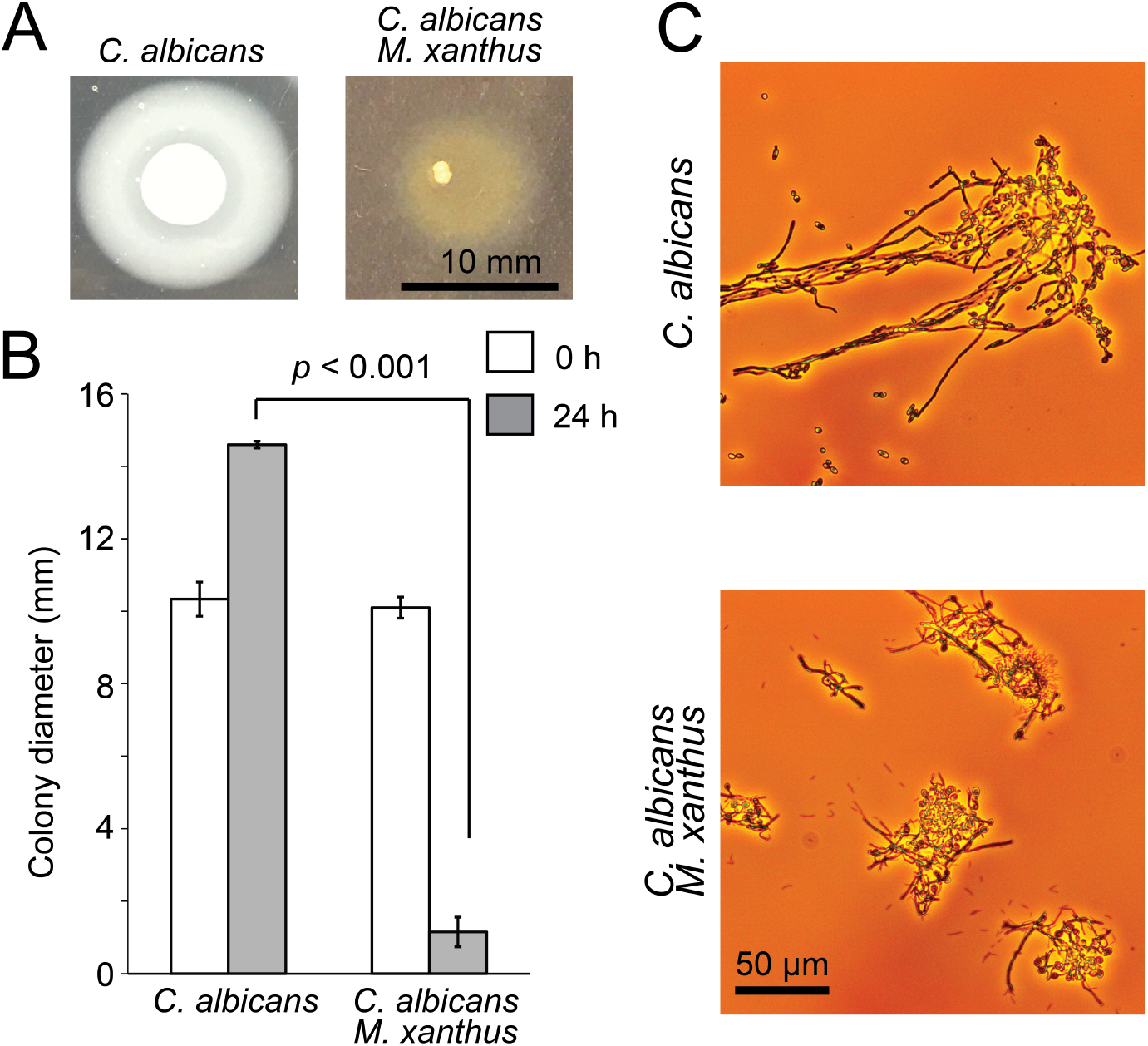
*M. xanthus* inhibits the growth and virulence of *C. albicans*. **A)** *M. xanthus* inhibits the growth of *C. albicans* on solid medium. **B)** Quantitative analysis of *C. albicans* colony expansion. Diameters were measured from the colonies of six biological replicates. **C)** *M. xanthus* inhibits hyphal growth of *C. albicans* in liquid medium.

## Materials and Methods

### Strains and growth conditions

*M. xanthus* DZ2, *C. neoformans var. grubii* strain H99 (serotype A) and *C. albicans* strain SC5314 were used as the wild-type strains. *M. xanthus* and fungal cells were cultivated at 30 °C using CYE (1% Casitone, 0.5% yeast extract, 10 mM 3-(N-morpholino) propanesulfonic acid (MOPS), pH 7.6, 4 mM MgSO_4_) and YPD (1% yeast extract, 2% Bacto Peptone, and 2% dextrose, 2% agar) medium, respectively. The formation of *C. neoformans* capsules and *C. albicans* hypha was visualized with a Nikon Eclipse™ 400 microscope and an OMAX™ A3590U camera.

### Dual-species interaction assays

To visualize the fungicidal activities of *M. xanthus* on solid surfaces, 150 μl (OD_600_ 0.2) *C. neoformans* cells were spread onto CYE plates that contain 1.5% agar, and air-dried. 6 μl of *M. xanthus* cells were spotted onto the plates pre-inoculated with fungal cells, air-dried, and incubated for 24 h in dark at room temperature. The clear zones on the *C. neoformans* lawn indicate growth inhibition.

The survival rates of *C. neoformans* and *M. xanthus* in liquid coculture were determined by CFU assays. *M. xanthus* and fungal cells were washed and resuspended in PBS. A mixture of fungal cells (500 μl, OD_600_ 1) and *M. xanthus* cells (500 μl, 4 × 10^9^ cfu ml^-1^) was co-incubated in a 24-well plate in 30°C shaker. After 24 h, 100 μl of diluted cell mixtures were plated onto YPD (with 100 μg/ml kanamycin) and CYE (with 0.1 mg/ml nourseothricin) plates and the CFUs of *C. neoformans* and *M. xanthus* were counted after a 48-h incubation at 30 °C.

To test the fungicidal activity of the filtered supernatant of liquid *M. xanthus* culture, *M. xanthus* was grown in liquid CYE medium at 32° C, till its OD_600_ reaches 1 (∼24 h). Then 10 ml supernatant was filtered (pore size 0.2 μm). To test the heat-resistance of the antifungal factors, when mentioned, the filtered supernatant was boiled at 100° C for 15 min. The supernatant was mixed with *C. neoformans* liquid culture (OD_600_ 1) at a ratio of 1:1. After 24-h of incubation at 32° C, 100 μl of series-diluted samples were plated onto YPD (with 100 μg/ml kanamycin) and viable cryptococcal cells were counted. For assessment of cell contact dependence, a two-chamber assay was performed as previously described^47^ with some modifications. *M. xanthus* (500 μl, 4 × 10^9^ cfu ml^-1^) or media only were inoculated into the bottom chamber of the 24-well plate containing a 0.4 μm membrane insert (? Check the manufacturer in the lab), *C. neoformans* (500μl, OD_600_ 1) was inoculated on the top chamber. At the same time, mixed *C. neoformans* and *M. xanthus* or *C. neoformans* and media without membrane insert were included as controls. At indicated time points, series of diluted samples were plated and viable cryptococcal cells were counted.

### Virtualization of *C. neoformans* melanin and capsule

*C. neoformans* melanin production was examined on solid L-3,4-dihydroxyphenyl-alanine (L-DOPA; 100 mg/l) medium. Overnight grown *C. neoformans* and *M. xanthus* cultures were washed, and a mixture of *C. neoformans* (3 μl, OD_600_ 1) and *M. xanthus* (3 μl, 4 × 10^9^ cfu ml^-1^) was spotted onto L-DOPA plates and incubated at 30°C in the dark for four days. Melanization was observed as *C. neoformans* colonies developed a dark brown color.

To measure the cryptococcal capsule formation, *C. neoformans* and *M. xanthus* cells were grown overnight in YPD and CYE medium, respectively and washed three times with PBS. Then *C. neoformans* cells (100 μl, OD_600_ 1) alone or mixed with *M. xanthus* (100 μl, 4 × 10^9^ cfu ml^-1^) were inoculated onto Dulbecco’s Modified Eagle Medium (DMEM) and grown for three days at 37°C. The thickness of *C. neoformans* capsules was measured using ImageJ.

### *C. albicans* hypha formation

*O*vernight *C. albicans* was washed twice in PBS and adjusted to OD_600_ 1. *M. xanthus* culture was washed twice and adjusted to series of concentrations (4 × 10^9^ cfu ml^-1^). Equal volumes of *C. albicans* and *M. xanthus* cells were mixed, and 6 μl of the mixture was spotted onto solid agar supplemented with 10% FBS. Pure *C. albicans* culture was included as a positive control. Plates were incubated at 30 °C for 24 h before being photographed.

### Fluorescence microscopy

Equal volumes of *C. neoformans* (100 μl, OD_600_ 1) and *M. xanthus* (100 μl, 4 × 10^9^ cfu ml^-1^) cells were mixed and grown at 30 °C for 6 h wish shaking. Cell mixtures were stained with 20 μM CFW and 5 μg ml^-1^ WGA before visualization. To visualize plasma membrane potential changes, samples were placed on a microscope slide with 0.8% agarose gel containing 5 μM Bis-(1,3-dibutylbarbituric acid) trimethine oxonol (DiBAC4(3)) (Biotium) and 0.75 μM permeability reporter PI (MP Biomedicals). To suppress binding of DiBAC4(3) on the glass surface, the coverslips were coated with L-dopamine^48^. Amphotericin B, used as a positive control to induce membrane depolarization without pore formation, was added to 3 μg ml^-1^. The pore-forming peptide mellitin was added to the cells 10 min before the end of the induction period at a final concentration of 10 μM^49^. Untreated *C. neoformans* monoculture was used as a control. Microscopy images were captured using a Hamamatsu ImagEM X2™ EM-CCD camera C9100-23B (pixel size 160 nm) on an inverted Nikon Eclipse-Ti microscope with a 1.49 NA TIRF objective. The fluorescence intensity was measured using ImageJ. Corresponding background values were subtracted from the measurements.

## Acknowledgements

We thank Dr. Xiaorong Lin for *C. neoformans* and *C. albicans* strains, Dr. Matthew Sachs for technical guidance. This work is supported by the National Institute of Health R01GM129000.

## Notes

### Competing Interest Statement

The authors have declared no competing interest.

